# Isolation and quantification of blood apoptotic bodies in neurological patients

**DOI:** 10.1101/757872

**Authors:** Gemma Serrano-Heras, Inmaculada Díaz-Maroto, Blanca Carrión, Ana B. Perona-Moratalla, Julia Gracia, Sandra Arteaga, Francisco Hernández-Fernández, Jorge García-García, Beatriz Castro-Robles, Oscar Ayo-Martín, Tomás Segura

**Affiliations:** Research Unit, Complejo Hospitalario Universitario de Albacete, Albacete, Spain; Department of Neurology, Complejo Hospitalario Universitario de Albacete, Albacete, Spain

**Keywords:** Extracellular vesicles, apoptotic bodies, plasma samples, biomarkers, stroke, neurodegenerative diseases

## Abstract

Dysregulation of apoptosis may contribute to the etiology and/or progression of several prevalent diseases, including stroke and neurodegenerative pathologies. So, detection of the apoptotic processes in patients would be useful in daily clinical practice, However, the *in vivo* analysis of apoptosis that occurs in tissue has limitations. We therefore propose to use circulating apoptotic bodies as biomarkers for measuring apoptotic death in patients. Since there is no scientific literature establishing the most appropriate method for measuring apoptotic bodies from human blood samples, we here describe a reproducible centrifugation-based method combined with electron microscopy, dynamic light scattering and flow cytometry studies to isolate, characterize and quantify plasma apoptotic bodies of patients with ischemic stroke, multiple sclerosis and Parkinson’s disease. The analysis revealed that our isolation protocol achieves notable recovery rates of highly-purified intact apoptotic bodies. This easy and rapid procedure would help physicians to implement the use of plasma apoptotic bodies analysis as a non-invasive tool to monitor apoptotic death in neurological patients for prognostic purposes and for following disease activity and assessing drug effectiveness.

## Introduction

Three major morphologies of cell death have been described: apoptosis (type I), cell death associated with autophagy (type II) and necrosis (type III)[1]. During apoptosis, decrease in cell volume, cytoplasmic condensation, mitochondrial membrane permeabilization, DNA fragmentation, and chromatin condensation followed by nuclear fragmentation and cytoplasmic membrane blebbing takes place, thereby leading to the total desintegration of the cell. In the late phase of apoptosis, and as a result of cell fragmentation, apoptotic bodies are generated [2,3]. These small membranous vesicles, which have been released into their microenviroment and blood circulation, are removed by phagocytosis thus avoiding the inflammatory response, in the same manner as described for apoptotic cells [4].

Recent studies have reported that besides apoptosis, programmed cell death also includes autophagic cell death [3–5]. This dying process involves the engulfment of cytoplasmic material and intracellular organelles within double-membrane vesicles, called autophagosomes and occurs without unequivocal morphological manifestations of apoptosis and formation of apoptotic bodies [6]. Likewise, the process of necrosis, traditionally considered as an unprogrammed death resulting from an overwhelming insult, is also morphologically distinct from apoptosis in many of its characteristics such as loss of membrane integrity and rapid cell and organelle swelling. Interestingly, increasing studies have described a genetically programmed and regulated form of necrosis, termed necroptosis. However, necroptotic death, like cellular necrosis, culminates in cell lysis and release of the cytoplasmic content into the surrounding tissue, provoking immediate inflammation [1,7,8].

It is now becoming evident that an increasing number of pathological situations can be related to aberrant apoptosis, such as cerebrovascular and neurodegenerative diseases [9–12]. Thus, the evaluation of the apoptotic processes, in these neurological diseases would be useful in daily clinical practice, since it would allow an earlier and more accurate prognosis, more effective monitoring of evolution of the disease, and the selection of the most appropriate treatment for each patient. However, the *in vivo* analysis of apoptosis that occurs in tissue has limitations as it is currently necessary either to obtain tissue samples of patients by interventionist methods, biopsy and surgery, or to inject fluorescent or radiolabeled probes into the whole body [13,14]. Therefore, there is an urgent medical need for non-invasive procedures that allow the study of apoptosis, preferably in a quick, simple and quantitative manner. In this context, the authors propose the determination of plasma levels of apoptotic bodies, as a means to measure in vivo apoptosis associated with cerebrovascular and neurodegenerative pathologies in a non-invasive and sensitive manner.

Hence, the aim of this study was to develop a brief time consuming, simple and reproducible method for isolating circulating apoptotic bodies that maintain structural integrity and can therefore be quantified. To address this, blood samples from patients with ischemic stroke, multiple sclerosis and Parkinson’s disease, pathological disorders in which the involvement of apoptosis has been postulated, were first collected and subjected to a protocol based on differential centrifugation. The resulting vesicle preparations were characterized by transmission electron microscopy (TEM) and dynamic light scattering analysis, and the DNA profile of these vesicles were also investigated using a Bioanalyzer. Next, flow cytometry was applied to determine the yield and purity accomplished during the developed method to isolate apoptotic bodies.

## Material and methods

### Patients and Blood sampling

This study was conducted with the informed consent of patients according to the Declaration of Helsinki, and the experimental protocol was approved by our institutional Human Ethics Committee. The cohort consisted of 16 patients, eight diagnosed with acute phase ischemic stroke, four diagnosed with multiple sclerosis and four with Parkinson’s disease. Detailed information about the patients is presented in Table 1. Peripheral blood (20 ml) from each patient was collected in the neurology service and deposited in tubes with sodium citrate. Within 2 hours post-extraction, whole blood was centrifuged at 160xg for 10 minutes at room temperature (RT) to separate plasma from blood cells (Figure 1 upper part). Plasma was transferred to a clean tube and stored for up to 3 days at 4°C until proceeding with apoptotic bodies isolation.

**Table 1.** Characteristics of the study patients

| **Patient code** | **Neurological disease** | **Gender** | **Age** | **Clinical features at time of blood collection** |
| --- | --- | --- | --- | --- |
| IS1 | Ischemic Stroke | Male | 82 | acute phase (undetermined etiology) |
| IS2 | Ischemic Stroke | Male | 55 | acute phase (undetermined etiology) |
| IS3 | Ischemic Stroke | Male | 72 | acute phase (cardioembolic etiology) |
| IS4 | Ischemic Stroke | Female | 78 | acute phase (undetermined etiology) |
| IS5 | Ischemic Stroke | Male | 57 | acute phase (cardioembolic etiology) |
| IS6 | Ischemic Stroke | Male | 79 | acute phase (undetermined etiology) |
| IS7 | Ischemic Stroke | Female | 84 | acute phase (cardioembolic etiology) |
| IS8 | Ischemic Stroke | Male | 64 | acute phase (cardioembolic etiology) |
| MS1 | Multiple sclerosis | Female | 37 | relapse episode (early RRMS* type) |
| MS2 | Multiple sclerosis | Female | 27 | relapse episode (early RRMS type) |
| MS3 | Multiple sclerosis | Male | 34 | relapse episode (early RRMS type) |
| MS4 | Multiple sclerosis | Female | 24 | relapse episode (early RRMS type) |
| PD1 | Parkinson's disease | Female | 65 | H&Y** stage II (tremor-dominant type) |
| PD2 | Parkinson's disease | Female | 62 | H&Y stage II (tremor-dominant type) |
| PD3 | Parkinson's disease | Female | 71 | H&Y stage II (akinetic-rigid type) |
| PD4 | Parkinson's disease | Female | 75 | H&Y stage II (akinetic-rigid type) |

Patient code, neurological disease, gender, age, clinical features at time of blood collection are shown.
\* RRMS: relapsing-remitting multiple sclerosis.
\*\* H&Y: Hoehn and Yahr scale.

**Figure 1.**
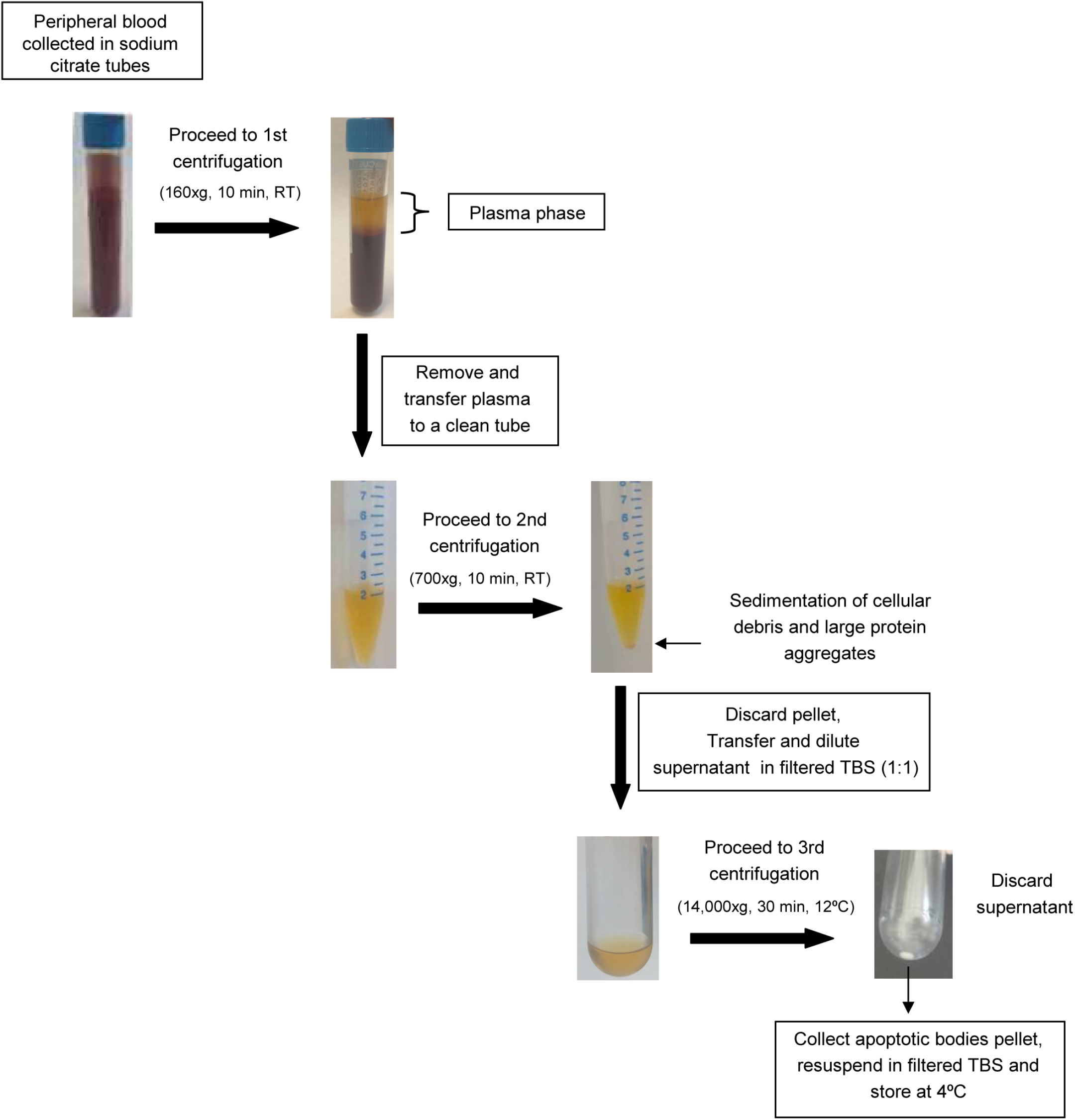
Flow chart of the designed protocol for isolating apoptotic bodies from blood samples. Whole blood was drawn from patients with either ischemic stroke or neurodegenerative pathology by venipuncture and collected into tubes that contained an anticoagulant (sodium citrate). The plasma phase were separated from blood cells by a first centrifugation (at 160xg for 10 minutes). After that, such plasma was centrifuged (at 700xg for 10 minutes) again to spin down the cellular debris. The fluid was then collected, diluted 1:1 with TBS 1X and further used to isolate apoptotic bodies. Apoptotic bodies were pelleted by centrifugation at 14,000xg for 30 minutes.

### Apoptotic bodies isolation from human plasma samples

Apoptotic bodies were isolated from plasma samples of patients with different neurological diseases, including ischemic stroke, multiple sclerosis and Parkinson’s, by differential centrifugation. The workflow for the preparation procedure of circulating apoptotic bodies is depicted in Figure 1. The plasma phase obtained from blood samples was centrifuged at 700xg for 10 minutes at RT to remove cellular debris and large protein aggregates. Then, the supernatant was transferred to a clean polycarbonate tube, diluted with an equal volume of Tris-buffered saline 1X (TBS)(50mM Tris-Cl, 150mM NaCl, pH:7.4), filtered with a 0.22 µm pore size hydrophilic Polyethersulfone (PES) membrane (Millipore Corporation, Bedford, MA), and centrifuged at 14,000xg for 30 minutes at 12°C to spin down apoptotic bodies (Optima L-100 XP ultracentrifuge, SW32 Ti rotor and thick wall Beckman tubes (Beckman Coulter)). Next, the remaining supernatant was eliminated and the pellet containing apoptotic bodies was resuspended by pipetting up and down gently in 0.22µm filtered-TBS 1X. The preparations of apoptotic bodies were immediately used for analysis or stored at 4°C for up to twelve months.

### Transmission electron microscopy examination

The sediments of apoptotic bodies obtained from 1ml of neurological patients plasma by the above-mentioned centrifugation-based protocol were submitted to electron microscopy study. Each pellet was resuspended in 300µl of the ice-cold fixation buffer (EM grade 2% glutaraldehyde (Sigma-Aldrich Chemical Co) in 1X TBS) and maintained on a mixing wheel for at least 24 hours at 4°C. After that, the samples were washed for 10 minutes with filtered Phosphate Buffer (PB)(0.1M, NaH2PO4 and Na2HPO4, pH:7.4) two times, and a single drop of 1.5 % agar was added. The embedded apoptotic bodies were post-fixed in 2% osmium tetroxide (OsO4) for 1 hour at RT and negatively stained with 2% uranyl acetate in the dark for 2 hours at 4°C. Then, the samples were rinsed in distilled water, dehydrated in ethanol, and infiltrated overnight in Durcupan resin (Fluka-Sigma-Aldrich, St. Louis, USA). Following resin polymerization, ultra-thin sections (0.08 µm) of apoptotic bodies were cut with a diamond knife, stained with lead citrate (Reynolds solution) and examined under a transmission electron microscope FEI Tecnai G2 Spirit (FEI Europe, Eindhoven, Netherlands). Images were taken using a digital camera, model Morada (Olympus Soft Image Solutions GmbH, Münster, Germany).

### Dynamic Light Scattering analysis

The isolated apoptotic bodies preparations were analyzed by dynamic light scattering (DLS) using a Malvern Zetasizer 5000 (Malvern Instruments Ltd, UK) to provide data on vesicular size distribution. DLS is an absolute sizing technique that determines size indirectly from measured speed of particles moving in suspension (larger particles travel more slowly, whereas smaller particles travel faster). When a sample consists of a mixture of molecules of different sizes, a polydispersity analysis is performed in order to determine the sizes profile of particles in the suspension as well as to assess the relative intensity (expressed as a percentage) of each particle with respect to all particles. In short, the apoptotic bodies samples were diluted 1:10 in 0.22 µm filtered PBS and placed in low volume disposable cuvettes for the analysis. Three measurements were carried out for each sample in which the DLS parameters (temperature, duration used, measurement position, dispersant refractive index and viscosity) remained unaltered within experiments. In all preparations, the median particle diameter size (nm), standard deviation, sizes distribution and the relative percentage of each small vesicle (relative intensity) were obtained using Zetasizer software

### Apoptotic DNA extraction and size determination by Bioanalyzer System

Prior to DNA isolation, the apoptotic bodies preparations were treated with 40µg/ml RNAse A (Promega Biotech Iberica S.L) and 27 KunitzU/ml DNAse I (Werfen-Qiagen N.V) for 30 min at 37°C, respectively, in order to remove possible contaminating external nucleic acids. After treatment, the enzymes were inactivated using RiboLock RNase Inhibitor (Fermentas) and heat inactivation. Next, total DNA was extracted from apoptotic bodies using QIAamp DNA Mini Kit (Werfen-Qiagen) following manufacturer recommendations. A 5µl-aliquot of extracted apoptotic DNA was used to determine quality and size distribution by electrophoretic separation on microfabricated chips using the Agilent 2100 Bioanalyzer System. These chips accommodate sample wells and become an integrated electrical circuit once the wells and channels are filled. Charged biomolecules like apoptotic DNA are electrophoretically driven by a voltage gradient, similar to slab gel electrophoresis. Because of a constant mass-to-charge ratio and the presence of a sieving polymer matrix, the target molecules are separated by size and subsequently detected via laser induced fluorescence detection. The software creates an electropherogram file as a graph representing fluorescence units (FU) against migration time, which is adjusted into DNA size using an internal size standard (a mixture of DNA fragments of known sizes; 25, 200, 500, 1000, 2000 and 4000 nucleotides).

### Apoptotic bodies quantification using Flow Cytometry

Over the last 10 years, to ease flow cytometry analysis of small molecules, vesicles have been coupled to beads that provide a larger surface and more scattered light [15]. However, this technical approach is inconvenient for subsequent use of vesicles for functional studies. Moreover, the detection could have imperfect reproducibility since it is largely dependent on the abundance/availability of antigen on such vesicles, which is recognized by the antibody coupled to the beads. In this way, the relative presence of various surface proteins can be determined, but this method would neither allow extracellular vesicle quantification nor discriminate between different vesicle subsets. Accordingly, we established a direct and multi-parameter quantitative method for flow cytometric analysis of isolated apoptotic bodies, without the necessity of using antibody-coated beads. The method relies on the fact that the membrane of apoptotic bodies, as that of apoptotic cells, is characterized by the presence of pores and phophatidylserine on the external surface [16]. Thus, staining with fluorophore-coupled Annexin V in conjunction with a vital dye, such as propidium iodide (PI), can identify and quantify apoptotic bodies due to the high affinity of Annexin V for phosphatidylserine and because the DNA-carrying apoptotic bodies with pores are permeable to such nucleic acid intercalating agent. Thereby, the amount of apoptotic bodies can be calculated as the number double positive events for Annexin V and PI (Annexin V^+^/PI^+^) recorded in the upper right area of the representative flow cytometry dot plot, and, subsequently, expressed as the number of isolated apoptotic bodies by the analyzed volume of the preparation. Meanwhile, the concentration of other small membrane impermeable vesicle types, including exosomes and microparticles, can be determined as single positive events for Annexin V and negative events for PI staining (Annexin V^+^/PI^−^). Specifically, 50µl of either isolated apoptotic bodies or plasma phase prior to the isolation procedure was diluted in 400µl of Annexin V binding buffer 1X (10mM Hepes/NaOH, 140mM NaCl, 2.5mM CaCl_2_;pH:7.4) and, subsequently, incubated with 40µl of propidium iodide (PI) dye (10mg/ml (Invitrogen) prepared in TBS 1X) overnight at 4°C in the dark. Following incubation, stained apoptotic bodies were labeled by adding 10µl of Annexin V conjugated with fluorochrome Dy-634 (Immunostep S.L.) for 2 hours at RT in the dark. Next, to remove unbound dyes, co-labeled apoptotic bodies were washed once with 2ml of TBS, centrifuged at 14,000g for 30 minutes at 12°C, resuspended in 500µl of Annexin V binding buffer 1X and, immediately, acquired at medium rate in a FACS Canto II flow cytometer (BD), which incorporates 2 air-cooled lasers at 488-and 633-nm wavelengths and the BD FACSDivaTM software. Forward (FSC-A) and side scatter (SSC-A) were measured on a logarithmic scale. The apoptotic body size gate was determined using “Calibration Beads”, which is a mix of size-calibrated fluorescent polystyrene beads with diameters of 0.22, 0.45, 0.88, 1.35, (Spherotech Inc) 3µm (Becton-Dickinson Biosciences). A similar strategy based on calibrated-beads has been previously proposed to determine the gate of microparticles, another type of small vesicle^19-21^. Specifically, in this study, the lower and upper limits of the apoptotic body gate were defined from just above the size distribution of 0.45 µm up to that of 3µm beads in a FSC-A (on a logarithmic-scale and a assigned voltage of 500Volts) and SSC-A settings (on a logarithmic-scale and a assigned voltage of 400Volts) with a fluorescence threshold set at 200Volts for fluorochrome Dy-634 parameter. Such discriminator was established in order to separate true events from background noise caused by Annexin V binding buffer. Furthermore, 2 staining controls; (i) 50µl of unstained apoptotic bodies diluted in Annexin V buffer and (ii) 50µl of Dy-634 labeled apoptotic bodies, were used to fix the level of background fluorescence (non-specific fluorescent signal). Thus, the events that appeared in this region (Apoptotic bodies gate) were further analyzed to monitor phosphatidylserine exposure and the presence of membrane pores and stained DNA using 660/20nm and 670LPnm fluorescence detection channels (on a logarithmic-scale) at a voltage of 350Volts and 470Volts, respectively. Hence, all samples were run with a medium flow rate of 90µl per minute and the number of double positive events for Dy-634 (absorption/emission max: 635/658nm) and PI (absorption/emission max: 535/617nm) was calculated. Measurements were made in duplicate and averaged.

## Results

### Morphological characterization and size distribution of the isolated vesicles preparations

Apoptotic bodies populations contained in the blood samples of a total of six patients, two diagnosed with ischemic stroke, two diagnosed with multiple sclerosis, and two diagnosed with Parkinson’s disease (clinical information for the patients is shown in Table 1), were collected following the described isolation protocol (Figure 1 and Material and methods) and, subsequently visualized by transmission electron microscopy (TEM) and analyzed by dynamic light scattering (DLS). Electron microscopic images of the isolated vesicle preparations showed round-shaped membrane structures containing compact, electron-dense chromatin distributed throughout the vesicles (Figure 2), which strongly resembles the described characteristics for apoptotic bodies isolated from cultured tissue and cells [17,18]. It is interesting to note that the microscopic examination revealed the presence of abundant intact vesicles whereas the finding of disrupted or fused vesicles was occasional, suggesting that the developed method based on serial centrifugations is able to isolate apoptotic bodies that preserve plasma membrane integrity. Next, the samples of isolated vesicles, morphologically similar to apoptotic bodies, were subjected to a light-scattering approach to obtain the size distribution of these apoptotic vesicles (Figure 3A). As depicted in the DLS analysis, the preparations displayed a homogeneous size distribution in which the main intensity particle population had a diameter ranging from 860 to 1350nm, from 800 to 1050nm and from 690 to 990nm for samples obtained of patients with ischemic stroke, multiple sclerosis and Parkinson’s disease, respectively. We additionally found, in all cases, a minority population, generally representing less than 20% of the total particles, with a size much smaller than that displayed by the majority vesicles. It suggests the presence, in the samples of isolated apoptotic bodies, of a limited proportion of other types of extracellular vesicles, such as microparticles and exosomes, which have been reported to be also present in human plasma.

**Figure 2.**
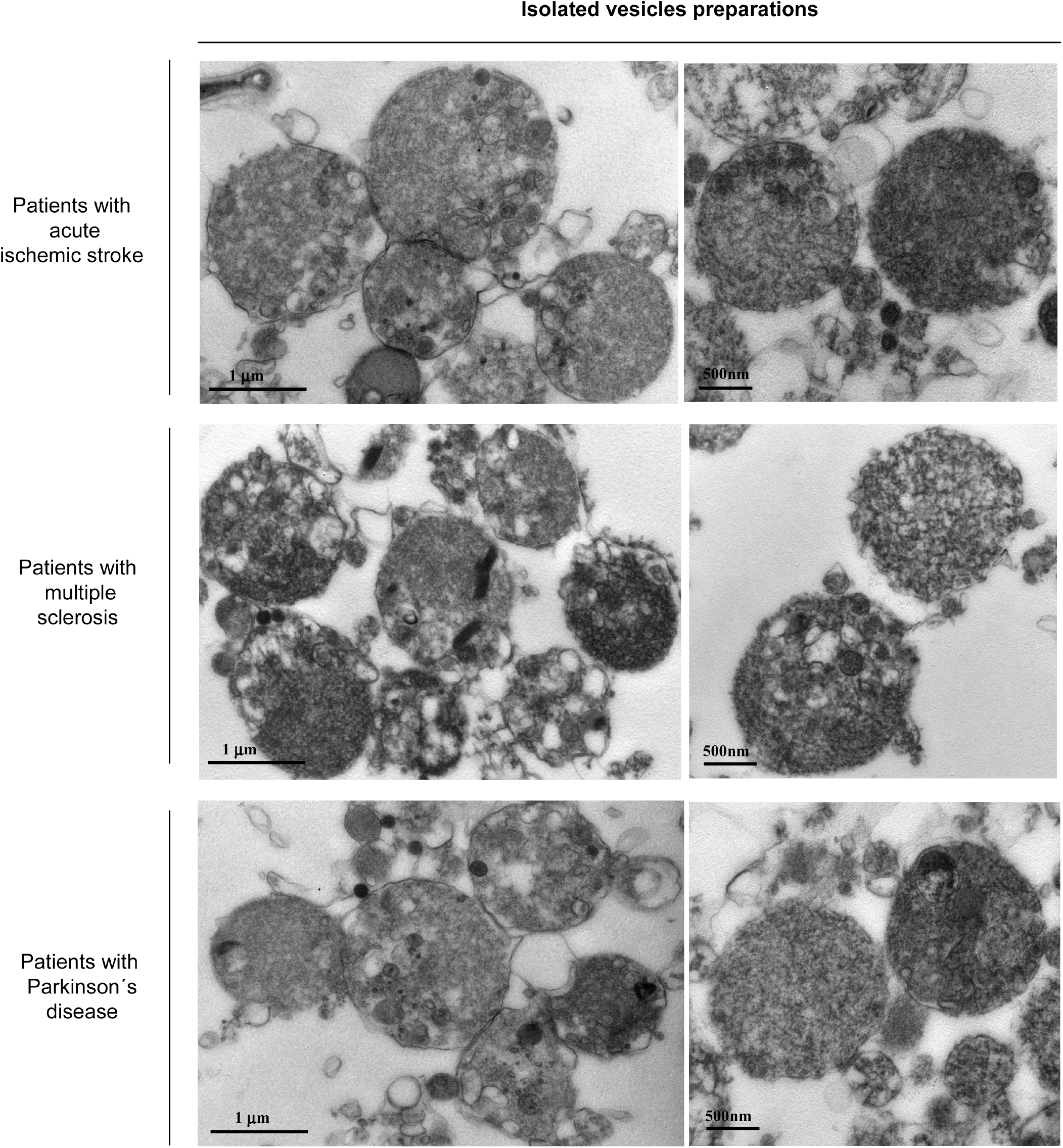
Analysis of the isolated vesicles by transmission electron microscopy. Representative electron micrographs of vesicles obtained from blood samples of stroke, multiple sclerosis and Parkinson’s disease patients following the described isolation protocol. All microscopy images show intact particles surrounded by a membrane, although some open membranes could be seen. Electron-dense chromatin substance appears within these rounded shaped vesicles, which have been found to display an average diameter of 1µm. The morphological features detected in most isolated vesicles are characteristic of apoptotic bodies, a type of extracellular vesicles. Scale bars; 1µm in the left panel, 500nm in the right panel.

**Figure 3.**
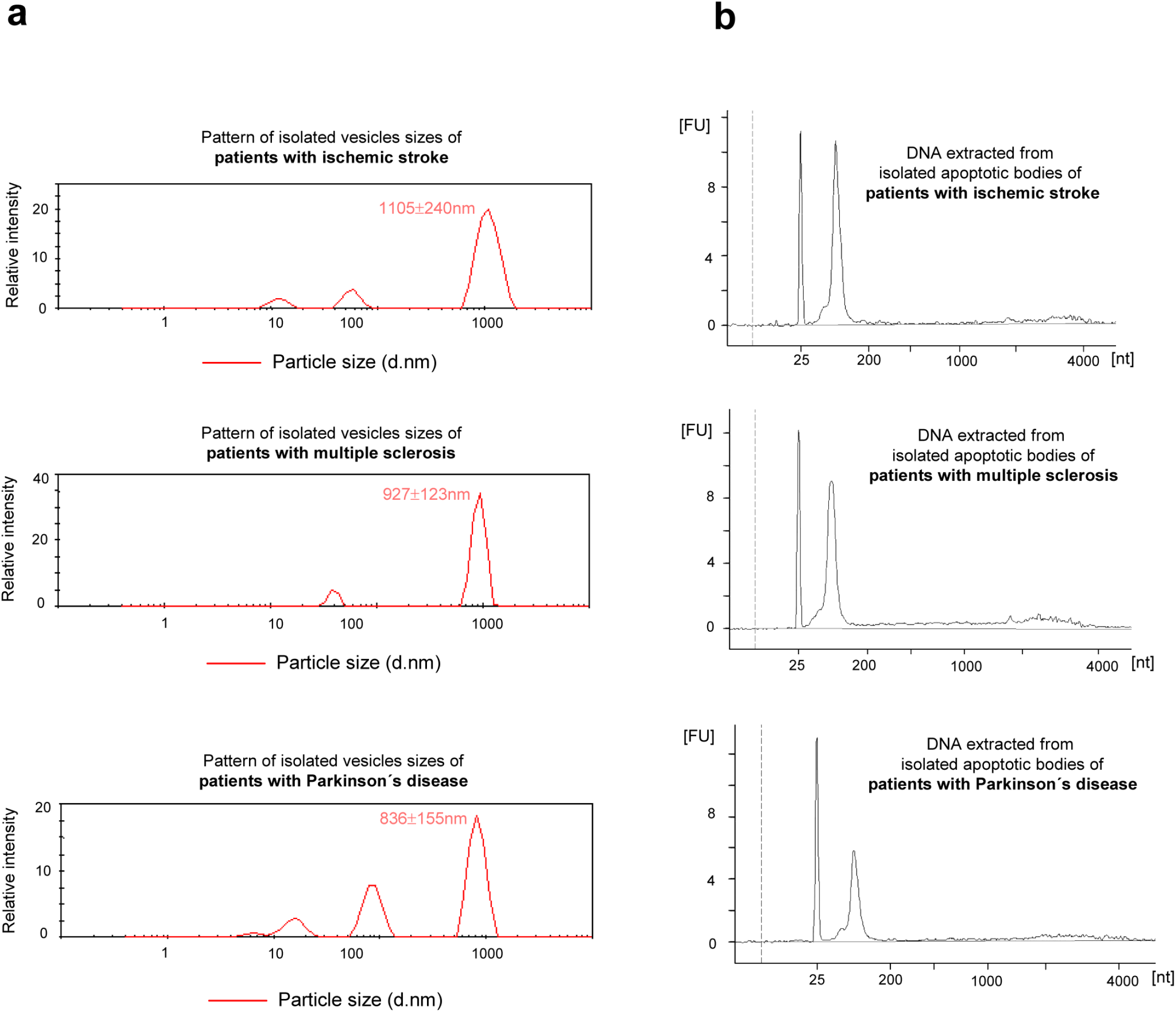
Vesicles size profiling and characterization of the vesicular DNA. **(A)** Size distribution from Dynamic Light Scattering analysis of plasma vesicles isolated from stroke, multiple sclerosis and Parkinson’s disease patients. Each graph is representative of three measurements and illustrates size (nm) versus intensity (relative frequency of each size range among the entire population of the isolated vesicles). **(B)** Size profile determination of the DNA extracted from patients plasma-isolated vesicles preparations using Bioanalyzer System. The electropherograms show sizes spectrum in nucleotides (nt) and fluorescence intensity (FU) of the DNA contained into the vesicles collected from neurological patients. Each individual graph represent one of three experiments with similar results.

### Nucleosome-sized DNA fragments packaged in plasma apoptotic bodies

Among the many markers of apoptosis, DNA cleavage (formation of 180-200bp DNA fragments) caused mainly by the action of nuclear endonucleases, is considered a hallmark [19]. Since, to date, it is unknown whether the DNA harbored by circulating apoptotic bodies has a similar pattern to that described in *in vitro* cells undergoing apoptosis, we further studied the DNA content of these vesicles using a Bioanalyzer, an automated on-chip electrophoresis system which is widely used for sizing of DNA [20]. Migration of DNA fragments extracted from patients with ischemic stroke, multiple sclerosis and Parkinson’s disease, respectively, was similar in the three gel matrices and there were no differences in the shape of the curves, regardless of the group of patient from which apoptotic bodies were isolated. Thus, the analysis revealed one sharp and dominant peak corresponding to a nearly 150-200bp target size and a weak almost negligible peak, indicating plasma apoptotic bodies exclusively carrying nucleosome-sized DNA fragments (Figure 3B).

### Assessment of yield achieved by the isolation apoptotic bodies method

In order to appraise the broad applicability to clinical routine workflow of our procedure for the isolation of human circulating apoptotic bodies, we next evaluated the isolation efficiency of such method by flow cytometry analysis. This technique offers the advantage of being commonly available at most research and clinical facilities and it appears reliable for the quantification and characterization of microvesicles [21–23]. In this study, we took advantage of three features of apoptotic bodies, external exposure of phosphatidylserine, presence of pores on the membrane and that they contain DNA [24], in order to analyze these vesicles by means of flow cytometry. Thus, the number of apoptotic bodies was measured as the number of double positive events in the representative flow cytometry dot plots, after incubation with fluorochrome-conjugated Annexin V, phosphatidylserine-binding protein, and propidium iodide (PI), a DNA intercalating agent that passes through membrane pores) (Figure 4A). Apoptotic bodies were previously defined by a gating method on the FSC-SSC dot plot using size-calibrated fluorescent beads.

**Figure 4.**
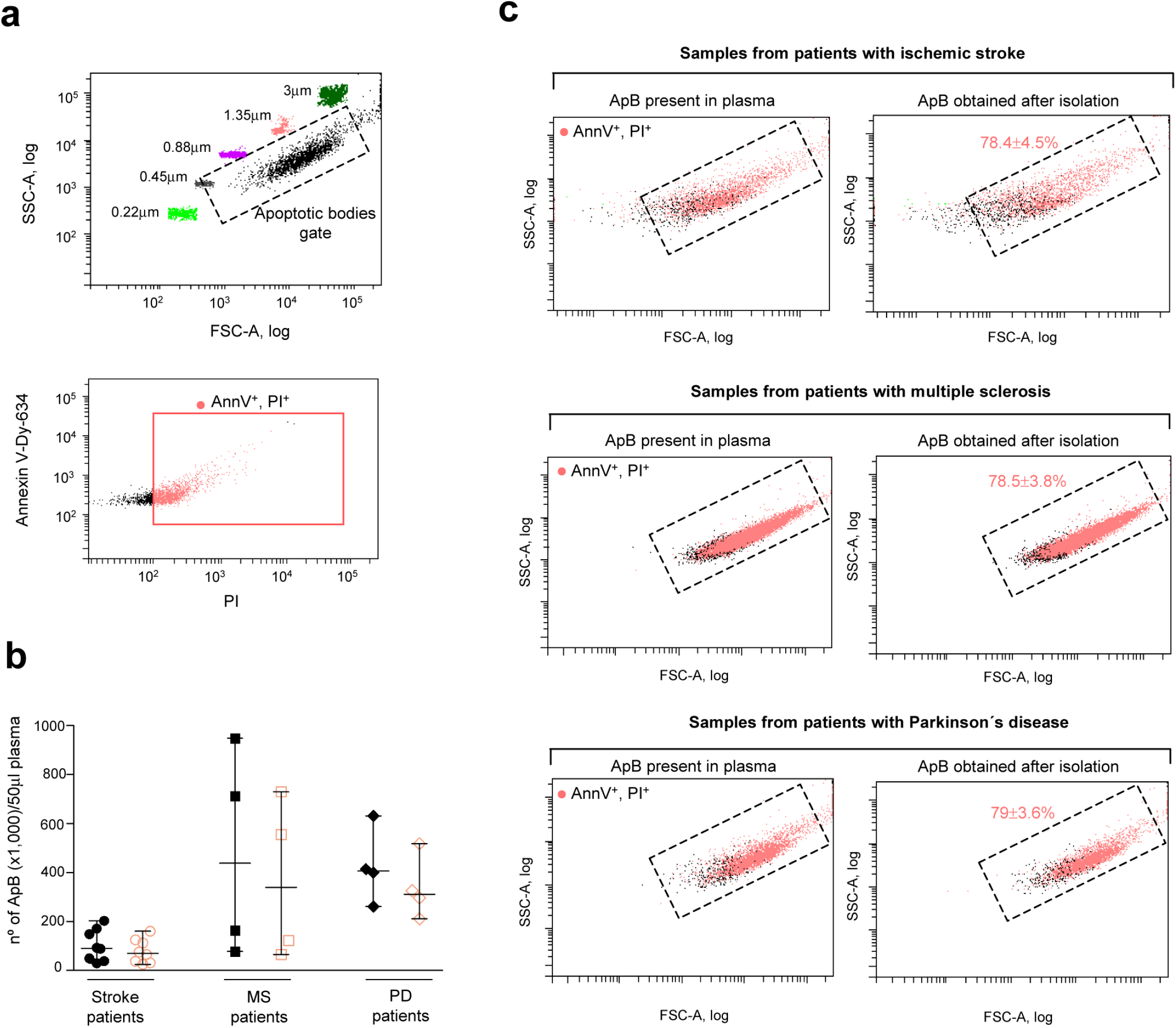
Selection strategy for the apoptotic bodies population and determination of the recovery efficiency on the isolation process. **(A)** Construction of the apoptotic bodies gate on the log-FSC-SSC plot using size calibrated-fluorescent beads are depicted (top). Representative flow cytometry plot of vesicles sample (previously gated on size selection) analyzed for the expression of apoptotic bodies markers using Annexin V-Dy-634/propidium iodide (PI) staining are shown. **(B)** The apoptotic bodies (ApB) concentration of the unprocessed plasma samples (filled black symbols) and the isolated vesicles preparations (empty pink symbols) from patients with ischemic stroke (*n*=8), multiple sclerosis (n=4) and Parkinson’s disease (n=4) are represented. **(C)** Representative analysis showing AnnexinV-Dy-634^+^PI^+^ apoptotic bodies (ApB) (pink points) on the FSC-SSC plots of plasma samples (left) and isolated vesicles (ApB) preparations (right) from neurological patients. The percentage yield of the isolation protocol, calculated by dividing the amount of isolated apoptotic bodies by the levels of these vesicles found in the starting plasma and multiply by 100, is indicated.

To calculate the yield of the isolation procedure, first, the concentration of isolated apoptotic bodies-sized vesicles in the preparations obtained from blood samples of 16 neurological patients was determined according to the number of AnnexinV/PI double positive events and the analyzed volume of each sample on the flow cytometer. Subsequently, such measurements were expressed as the number of isolated apoptotic bodies by a 50µl-fixed volume of initial plasma (Figure 4B). In parallel, in each subject blood sample, before proceeding to isolate apoptotic bodies, one part of the plasma phase was removed, stained with APC-Annexin V and PI, and analyzed, in the same manner, using the flow cytometer to calculate the total number of apoptotic bodies present in a 50µl-fixed volume of the unprocessed plasma (Figure 4B). Next, the comparison of the amount of apoptotic bodies detected by flow cytometry in the preparations obtained after the isolation method and the levels of these apoptotic vesicles found in the unprocessed plasma allowed us to verify that a high abundance of apoptotic bodies was sedimented after applying the developed centrifugation-based procedure. Specifically, the pellets obtained from eight patients with acute ischemic stroke, four patients with multiple sclerosis and four patients with Parkinson’s disease contained 78.4%±4.5%, 78.5%±3.8% and 79%±3.6%, respectively, of the total apoptotic bodies pool present in the original plasma (Figure 4B,C). These findings clearly indicate that our procedure efficiently pellets down a substantial amount of the apoptotic bodies present in blood samples. The high throughput recorded was reproducible in all neurological patients, regardless of their disease.

### Determination of purity degree of the isolated apoptotic bodies samples

For extracellular vesicle research, it is not only important that the isolation methodology provides high recovery rates but also that it is further accompanied by a low degree of other microvesicle contamination. Therefore, we evaluated the purity of the isolated apoptotic bodies preparations obtained from blood samples of the aforementioned neurological patients. To this end, the annexinV/PI stained isolated vesicles preparations were similarly analyzed by flow cytometry with the additional aim of quantifying all extracellular membrane vesicles present in such samples. Given that the diversity of secreted vesicle subpopulations, including apoptotic bodies and other smaller vesicles, has been reported as characterized by phophatidylserine externalization on cell membranes [25,26], first, the concentration of the membrane vesicles pool in the isolated apoptotic bodies samples was calculated as the total number of positive events for Annexin V by a 50µl-fixed volume of the preparation. Next, the amount of apoptotic bodies in the samples was measured as described previously as double-positive events for Annexin V and propidium iodide, but in relation to a 50µl-fixed volume of the isolated vesicles sample. Finally, the number of non-apoptotic vesicles by a 50µl-fixed volume of the preparation was further determined on the basis that these particles are recorded as positive events for Annexin V and negative for PI (Figure 5A). The relative percentage of apoptotic bodies in the preparations obtained from stroke patients as well as in the samples collected from subjects with neurodegenerative diseases was more than 80% while the amount of microvesicles and exosomes found comprised less than 20% of total vesicles (Figure 5B). Collectively, the flow cytometry studies demonstrate that the developed method is capable both of producing considerable recovery rates in the obtaining circulating apoptotic bodies and of successfully removing (up to 80%) plasma vesicle contaminants.

**Figure 5.**
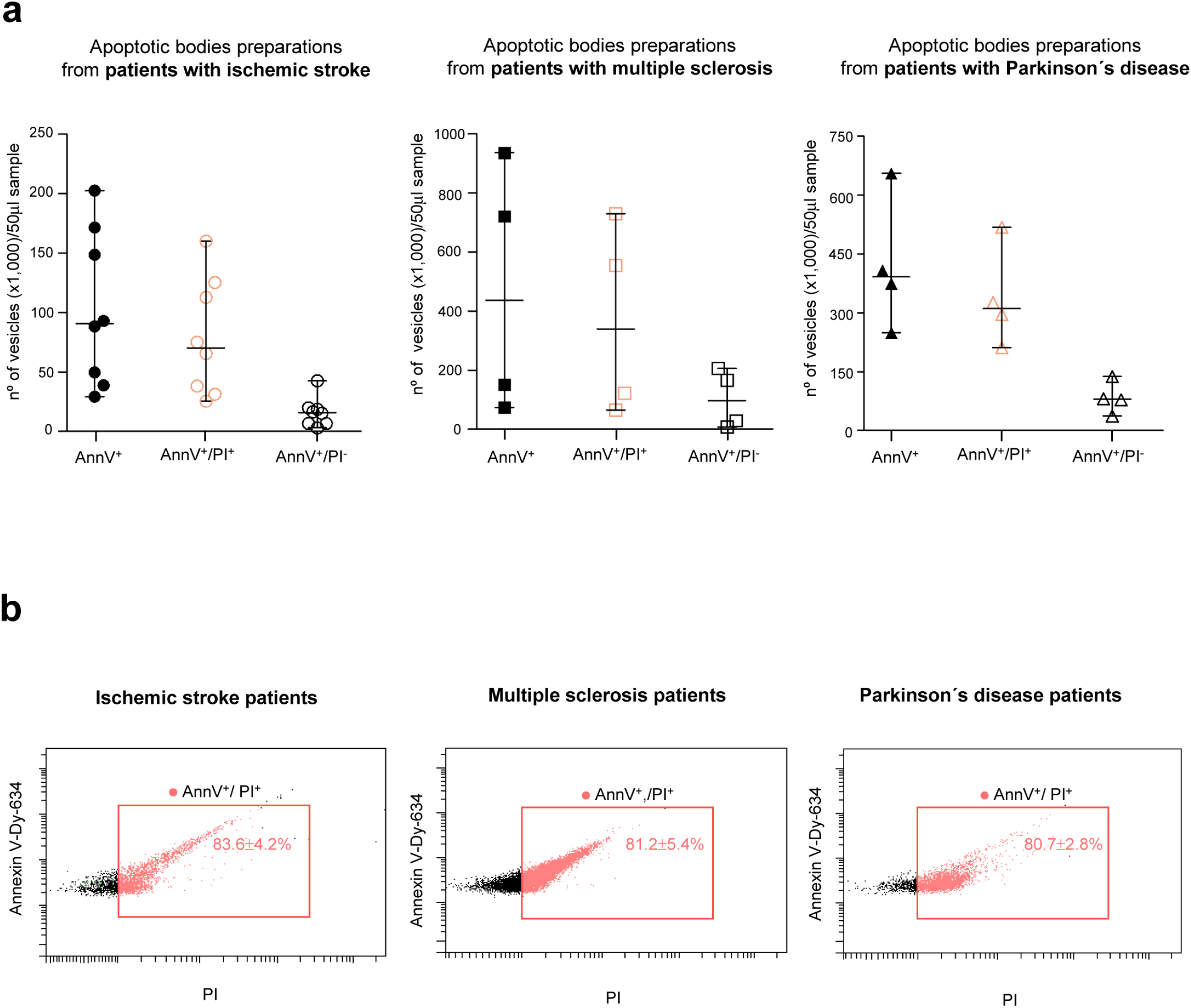
Assessment of the purity level on the apoptotic bodies isolation method. **(A)** The dots graphs show the flow-cytometry data analysis of isolated apoptotic bodies preparations from a total of 16 patients, eight diagnosed with ischemic stroke (left), four diagnosed with multiple sclerosis (middle) and four patients with Parkinson’s disease (right). The amounts of the total membrane vesicles, stained positively with Annexin V (AnnV) (filled black symbols), of the apoptotic bodies population, measured as double positive counts for annexin V and PI (empty pink symbols), and of the non-apoptotic vesicles population showing a annexin V-positive and a PI-negative signal (empty black symbols), expressed as the absolute number of each vesicles subsets counted in 50µl-volume of the apoptotic bodies fractions, are represented. **(B)**Representative flow cytometry profiles of signal intensity in dual staining for Annexin V and propidium iodide of the apoptotic bodies preparations from neurological patients. The relative percentage of gated AnnexinV^+^PI^+^ apoptotic bodies found in the isolated vesicles samples is indicated.

## Discussion

Extracellular vesicles (EVs), heterogeneous membrane vesicles released by cells into their microenvironment and blood circulation, are considered to play a fundamental role in many physiological and pathological processes [27–29]. Apoptotic bodies, the largest EVs (800-5000nm in diameter), are released by all cell types during the late stages of apoptosis. Microparticles or microvesicles, defined as 200-1000nm in size, are formed from the outward blebbing of the plasma membrane and subsequent shedding into extracellular space, whereas exosomes, the smallest membrane vesicles (40 and 100nm in diameter), are liberated into extracellular space by the fusion of the multivesicular body with the plasma membrane [30]. Research into EVs has been rising during the last decade, being microvesicles and exosomes the best characterized vesicle populations [25]. Indeed, there is available and updated information elucidating the most efficient methods for obtaining high yields of these highly-purified vesicles from both cell culture and complex biological fluids, such as plasma [31–33]. However, it is striking that apoptotic bodies have often been overlooked in studies of circulating vesicles. To date, no procedures for isolating apoptotic bodies from blood samples in which the percentage recovery and grade of contamination with other vesicles have been detailed. In this context, the authors here report an easy-to-perform and rapid protocol for apoptotic body collection from neurological patient-derived plasma samples. The microscopic examinations along with dynamic light scattering analysis revealed that the circulating vesicles isolated from acute stroke patients and subjects suffering multiple sclerosis and Parkinson’s disease, displayed morphology and sizes corresponding to apoptotic bodies. More importantly, these studies demonstrate that, using the developed protocol, it is possible to recover circulating intact apoptotic bodies that could be quantified using flow cytometry techniques, in order to assess the yield and purity obtained by the proposed isolation method. Interestingly, the results revealed that such procedure is able to collect the majority of apoptotic bodies present in the plasma of patients with ischemic stroke and neurodegenerative diseases. Furthermore, the isolated apoptotic bodies exhibited a high level of purity, since the proportion other microparticles was less that 20%. It is interesting to mention that another relevant and practical application of getting isolated apoptotic bodies that maintain the integrity of their membrane is that, in addition to its quantification, the identification of the cell type from which they are generated as a result of cell death via apoptosis is achievable, by the use of antibodies against a cell type-specific surface marker.

It should be highlighted that the isolation procedure of unbroken apoptotic bodies from neurological patients blood samples could be of great utility for *in vivo* study of apoptosis in pathologies, such as cerebral ischemia, multiple sclerosis and Parkinson’s disease, in which alteration of such cell death type has been identified as the main mechanism involved [9–12]. Specifically, the quantification of circulating isolated apoptotic bodies would potentially serve as a non-invasive tool for determining the degree of apoptotic cell death that has occurred in a particular neurological patient, which is, in fact, highly relevant to the understanding and treatment of such human diseases.

Diverse precise methods have been developed to examine the extent of apoptosis, among which the most widely used is the TUNEL assay, relies on the determination of DNA fragmentation and the detection of phosphatidylserine on the outside of plasma membrane of apoptotic cells using Annexin V [13]. These conventional approaches, however, include tissue sampling using invasive techniques, which makes it difficult and challenging for implementation in daily clinical practice. To overcome this limitation, several techniques for non-invasively detection and imaging of apoptosis have been developed and assayed in mouse models as well as in patients over the last years. These molecular imaging studies have mainly focused on the use of fluorescent or radiolabeled annexin V, protein that binds selectively to phosphatidylserine, a recognition signal on the outer cell membrane of apoptotic cells [34]. However, these detection approaches presents numerous challenges, including the presence of high levels of background fluorescence due to the presence of numerous fluorescent species in biological samples tissues [35] and the massive accumulation of the radionuclide in the kidney, spleen and liver that limit the amount of activity that can be administered and, hence, the sensitivity of the approach [36]. Furthermore, externalization of phosphatidylserines does not exclusively occur during apoptosis but also during other pathological conditions, such as under high-stress situations [37]. Another strategy in apoptosis imaging tries to target intracellular caspases, a key player in the apoptotic cascade. Development of protease-specific probes allows monitoring of caspase activation in vivo by a reporter gene assay, in which luciferase is activated upon cleavage of a caspase3-specific DEVD peptide motif [38]. Nonetheless, this approach, based on the use of a bioluminescent probe, cannot be directly translated into the clinic due to the need for intracellular delivery of reporter construct or to have luciferase-expressing cell lines.

In this context, the authors describe in this work a new mean of monitoring apoptotic death in living that presents an alternative to address the inherent challenges of apoptosis detection shown by aforementioned other methods. The analysis of circulating apoptotic bodies levels would eliminate both the need for extensive tissue collection, presented by the conventional approaches, and the requirement of the administration of fluorescent/radiolabeled compounds or reporter gene, as occurs for the apoptosis imaging techniques. More precisely, determination of plasma apoptotic bodies could be used as a novel prognostic factor and, furthermore, as an innovative marker for monitoring and evaluating the effectiveness of a given therapeutic treatment in pathologies associated with dysregulated apoptosis. For instance, in the context of neurodegenerative and cerebrovascular pathologies, lower levels of apoptotic bodies detected in the subsequent blood sample would reveal that the proven therapy is working. Very importantly, the ability to analyze apoptosis noninvasively and dynamically over time using plama apoptotic bodies would provide an opportunity for *in vivo* high-throughput screening of proapoptotic and antiapoptotic compounds. Alternatively, it should be noted that isolation and quantification of circulating apoptotic bodies could be a tool to take into account for providing new insights into the pathogenesis of specific diseases, which currently remain unclear. Recent studies have suggested that alterations in the core components of necrosis or a defective autophagy pathway could be other pathological mechanisms that either acts alone or in conjunction with apoptosis as a trigger to progression of some pathologies including certain neurodegenerative diseases, lysosomal storage disorders and several cancers [5,8,10]. Therefore, to better understand unwarranted loss of cells in human diseases of unknown etiology, it would be important to assess whether and to what extent apoptotic death is implicated. For this purpose, the accurate measurement of apoptotic bodies present in blood samples would be of great value.

In conclusion, the isolation and quantification of plasma apoptotic bodies represents an important new tool in the continuously growing approaches for the in vivo apoptosis detection, with significant improvement to tried-and-tested and experimental available research techniques. These apoptotic vesicles may have potential to serve as a valuable disease biomarker in the prognosis and evaluation of therapeutic efficiency in diseases such as ischemic stroke, multiple sclerosis and Parkinson’s disease. Nonetheless, due to the lack of standardized purification methods, the scientific community has not yet fully taken advantage of such apoptotic bodies. In this line, the authors have elaborated a minimally time-consuming and easy to achieve method for high yield isolation of highly purified intact apoptotic bodies from neurological patients blood samples. Our procedure is easily reproducible, of low economic cost and uses common laboratory ware and equipment, which facilitate its immediate application in a daily hospital routine.

## Abbreviations

DLS: dynamic light scattering
EV: extracellular vesicles
PI: propidium iodide
TEM: transmission electron microscopy
TUNEL: terminal deoxynucleotidyl transferase dUTP nick end labeling

## Acknowledgements

The patients participating in the study are gratefully acknowledged.

Authors thank Mario Soriano, technician of the Electron Microscopy Service in the Centro de Investigación Principe Felipe, Spain, and Esther Masiá and Inmaculada Conejos-Sánchez for their excellent technical assistance in the Dynamic Light Scattering studies. The Bioanalyzer analysis was provided by The Genomics and NGS Core Facility at the Centro de Biología Molecular Severo Ochoa,Spain.

This research was supported by a grant (number:PI12/01544) from the Instituto de Salud Carlos III (Spanish Ministry of Economy and Competitiveness) and co-financed by European Regional Development Fund (FEDER)

